# Nicotine and Opioid Polysubstance Abuse: Enhancement of opioid self-administration by systemic nicotine and modulation of opioid-associated memories by insular nicotine

**DOI:** 10.1101/2020.09.07.286542

**Authors:** Gregory C Loney, Christopher P King, Paul J Meyer

**Affiliations:** Program in Behavioral Neuroscience, Department of Psychology, The State University of New York, University at Buffalo, Buffalo, NY

**Keywords:** Contextual-Conditioning, Nicotine, Opioids, Polysubstance, Self-administration

## Abstract

Concurrent nicotine use is associated with increased liability for the development and exacerbation of opioid-use disorders. Habitual use of nicotine containing products increases propensity to misuse prescription opioids and its prevalence is substantially increased in individuals currently involved in opioid-treatment programs. Nicotine enhances self-administration of many classes of drugs in rodents, though evidence for direct effects on opioids is lacking. We sought to measure the effects of nicotine pretreatment on the reinforcing efficacy of opioids in both self-administration and contextual conditioning paradigms. First, we measured the effect of systemic nicotine pretreatment on self-administration of two opioids. Additionally, we measured the degree to which systemic nicotine pretreatment impacts the formation of morphine-associated contextual memories in conditioned taste avoidance and place preference paradigms. Given the involvement of the insula in the maintenance of substance abuse, its importance in nicotine addiction, and findings that insular inactivation impairs contextual drug conditioning, we examined whether nicotine administered directly to the insula could recapitulate the effects of systemic nicotine. We demonstrate that systemic nicotine pretreatment significantly enhances opioid self-administration and alters contextual conditioning. Furthermore, intra-insula nicotine similarly altered morphine contextual conditioning by blocking the formation of taste avoidance at all three morphine doses tested (5.0, 10, & 20 mg/kg), while shifting the dose-response curve of morphine in the place preference paradigm rightward. In conclusion, these data demonstrate that nicotine facilitates opioid intake and is partly acting within the insular cortex to obfuscate aversive opiate memories while potentiating approach to morphine-associated stimuli at higher doses.

## Introduction

Concurrent nicotine use is significantly correlated with substance abuse across a wide variety of commonly abused drugs, including opioids ^1-5^. Approximately 83-95% of individuals currently involved in an opioid-treatment program (OTP) are current users of nicotine and a positive relationship exists between nicotine use and propensity to misuse prescribed opioids ^6-8^. Because initiation of nicotine use typically precedes illicit drug use ^9^, nicotine-containing products may either promote the development of opioid-use disorders (OUDs) or exacerbate their severity. There is emerging evidence that nicotinic and opioid receptor systems interact in a manner that alter the response to opioid-associated cues in a fashion that may promote vulnerability to OUDs ^10-13^. Specific mechanisms and brain areas involved in these effects of nicotine in preclinical models have not been fully elucidated. Furthermore, while concurrent nicotine use increases methadone consumption in clinical populations ^14^ there are minimal reports of direct effects of nicotine pretreatment on opioid self-administration in preclinical models.

The formation and recall of drug-associated memories is indispensable for the development and maintenance of substance use disorders (SUDs). Mnemonic representations of the past reinforcing and aversive properties of drug administration and withdrawal can be elicited by cues and contexts that were present during drug episodes and can serve to elicit either approach or avoidance ^15-18^. Importantly, there is ample evidence for nicotine in modulating the development, strength, and recall of drug-associated memories across multiple classes of abused drugs and across multiple learning paradigms designed to measure the impact of the reinforcing and aversive effects of drug administration ^19,20 21-23^. Specifically, acute nicotine pretreatment robustly increases the incentive properties of discrete drug-predictive cues ^19,24-26^ and this enhancement is likely dependent on nicotinic activity within the midbrain dopaminergic circuitry, particularly the ventral tegmental area ^27,28^. There is emerging evidence that, unlike the effects seen on discrete stimuli, nicotine pretreatment may impair the strength of drug-associated contextual memories. Acute systemic and intracranial nicotine pretreatment limits the acquisition of conditioned place preference (CPP) to lower doses of morphine and cocaine^11,29,30^ and conditioned place avoidance from naloxone-precipitated morphine withdrawal ^31,32^. Additionally, across multiple commonly abused substances, acute systemic nicotine pretreatment impairs the formation of conditioned taste avoidance (CTA) following contextual presentation of a taste stimulus that predicts passive administration of the drug stimulus ^12,22,23^. Interestingly, at least for ethanol, discrete presentation of the taste stimulus, wherein the taste is contingently yoked to drug administration, results in a conditioned preference ^33-35^ further suggesting that the circumstances of presentation of a specific cue affect its associative properties ^36^ and thus potentially the relative contribution of potential brain areas and neurotransmitter systems.

One brain area in which nicotine may act to impair the formation of drug-associated contextual memories is the insular cortex (IC). The IC is a heterogeneous cortical structure that sends and receives numerous projections associated with sensory information and limbic processing and as such represents an area critical for integration of sensory and motivational functions ^37^. As a whole, the IC is critical for the acquisition and expression of both reinforcing and aversive drug-associated contextual memories and is heavily implicated in nicotine addiction ^38,39^. Inactivation of many IC subareas can affect the self-administration of commonly abused drugs ^40-42^, the strength of drug-induced CTA ^43^, as well as the strength of both the CPP ^44^ induced by drugs and the conditioned place avoidance induced by drug withdrawal ^45^. Activation of nAchRs on IC GABAergic interneurons facilitates synaptic depression at layer V pyramidal neurons, the primary excitatory output of the IC to subcortical structures ^46-48^. Morphine regulates the activity of inhibitory interneurons within the frontal cortex and these actions on interneurons appear to be required for contextual place conditioning ^49^. As such, nicotine pretreatment within the IC may have functional consequences on the formation and/or strength of opioid-associated memories. Specifically, the impairment of synaptic potentiation within the insular cortex by nicotinic-modulation of GABAergic interneurons may result in a deficit in learning about drug-associated contexts in a manner that promotes the development of SUDs.

Given the large overlap in nicotine and opioid dependence in clinical populations and the relative paucity of experimental data on the effects of nicotine on self-administration of opioids in preclinical models we examined the effects of nicotine on opioid intake and contextual conditioning. Furthermore, we focused on the insular cortex because it is critical for learning drug-associated contexts, is involved in the maintenance and development of substance abuse, is critical for the abuse-related effects of nicotine addiction ^40,50^, and insular plasticity appears to be modified through application of nicotine. First, we sought to demonstrate that nicotine pretreatment has functional consequences on opioid self-administration by examining whether it enhanced intake of the ultra short-acting synthetic opioid remifentanil in a fixed-ratio intravenous self-administration paradigm (IVSA). We also tested the effects of nicotine on a longer acting opiate in morphine. Next we examined the effects of systemic nicotine pretreatment on contextual conditioning by the interoceptive properties of multiple doses of morphine in CTA and CPP paradigms. Finally, we aimed to examine whether intra-insular delivery of nicotine similarly altered contextual conditioning with morphine as would be predicted by recent *in vitro* and behavioral studies ^11,21,46^.

## Results

### Experiment 1

Relative to responding under saline conditions, pre-treatment with nicotine significantly increased the number of active lever presses (*F*_(1,8)_ = 20.83, *P* < 0.01) resulting in an increased number of remifentanil infusions (*F*_(1,8)_ = 22.19, *P* < 0.01) while having no effect on the number of inactive lever presses (*P* = 0.70; Fig 1A). Analysis of the cumulative intake of remifentanil across the two-hour session grouped into 5-minute bins revealed that nicotine pretreatment, relative to saline, significantly elevated remifentanil intake beginning at 40 minutes and remained elevated throughout the remainder of the two-hour session (*F*_(23,184)_ = 26.00, *P* < 0.001; Fig 1B). Similarly, nicotine pretreatment significantly enhanced active responding for morphine (*F*_(1,8)_ = 12.77, *P* < 0.01) with no effect on inactive responding (*P* = 0.52) leading to an overall enhancement of morphine intake (*F*_(1,8)_ = 12.76, *P* < 0.01; Fig 1C). Analysis of cumulative morphine intake revealed that nicotine increased morphine intake beginning at 10 minutes and remained elevated throughout the session (*F*_(23,184)_ = 4.32, *P* < 0.001; Fig 1D).

**Figure 1 -.**
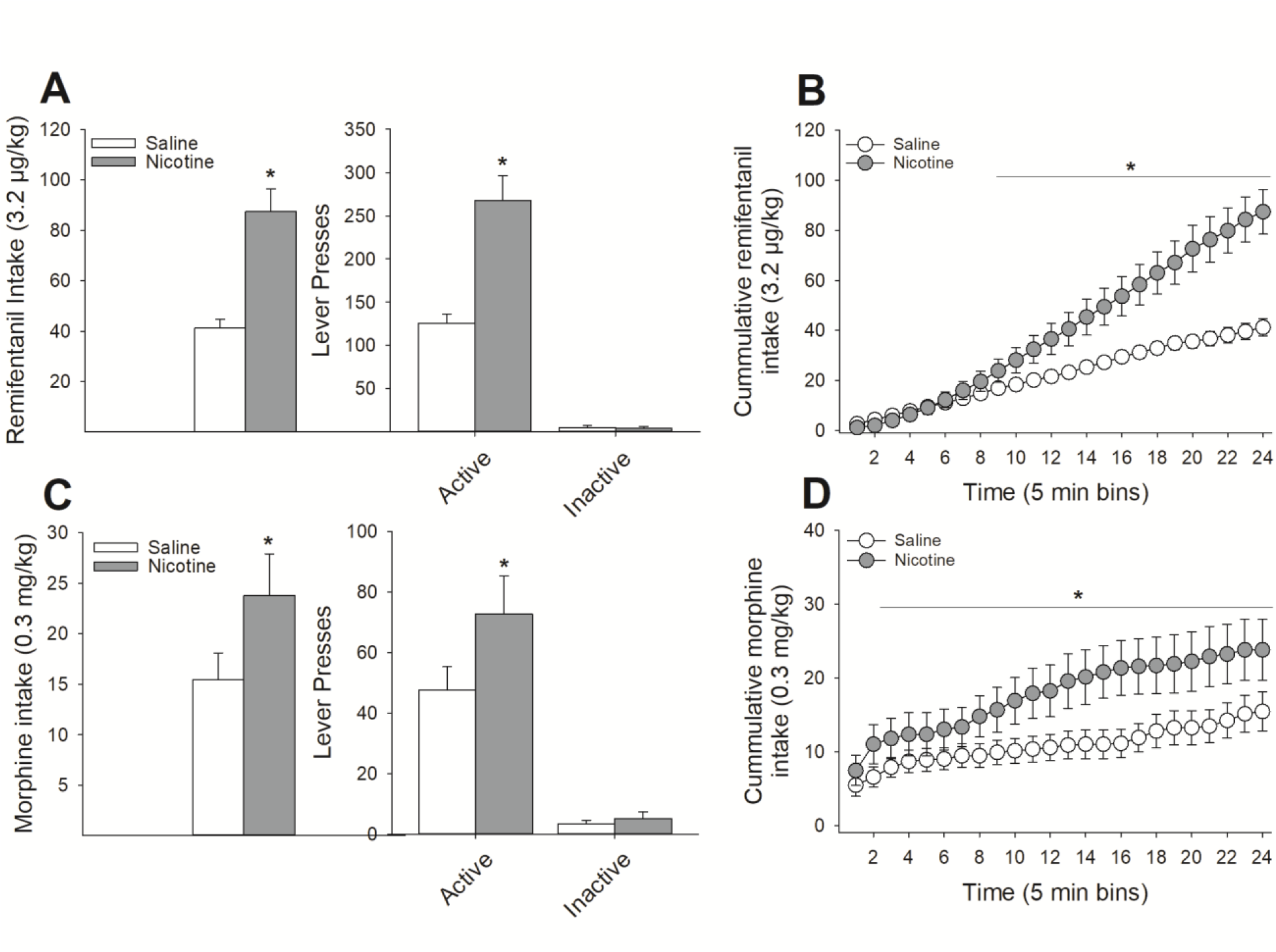
Systemic nicotine pretreatment enhances opioid self-administration. (**A**) In a within-subjects design, pretreatment with 0.4 mg/kg nicotine significantly increased the number of remifentanil infusions relative to saline pretreatment with no impact on responding on the inactive lever. (**B**) Time course analysis revealed that nicotine significantly increased remifentanil intake around the 40^th^ minute of the session. Nearly identical effects were observed with regards to nicotine pretreatment on morphine intake, with significant nicotine-induced increases in intake (**C**) such that nicotine increased morphine intake at the 10^th^ minute of the session and remained elevated throughout the session (**D**). *s indicate significant group differences (*P* < 0.05).

### Experiment 2

#### Conditioned Taste Avoidance

Systemic nicotine pretreatment (0.4 mg/kg) significantly attenuated the development of CTA to all doses of morphine tested (5.0, 10, & 20 mg/kg), relative to saline pretreatment (Fig 3a & b). A Drug x Dose x Day ANOVA conducted on the intake of the CS+ during the acquisition phase revealed a significant 3-way interaction (*F*_(9,192)_ = 2.40, *P* < 0.05). Post-hoc analyses indicated that nicotine-treated rats consumed significantly more sucrose than saline-treated rats on day 4 of conditioning with 5.0 mg/kg morphine, days 3 and 4 of conditioning with 10 mg/kg morphine, and days 2, 3, and 4 of conditioning with 20 mg/kg morphine (*P*s < 0.05; Fig 3a).

**Figure 3 -.**
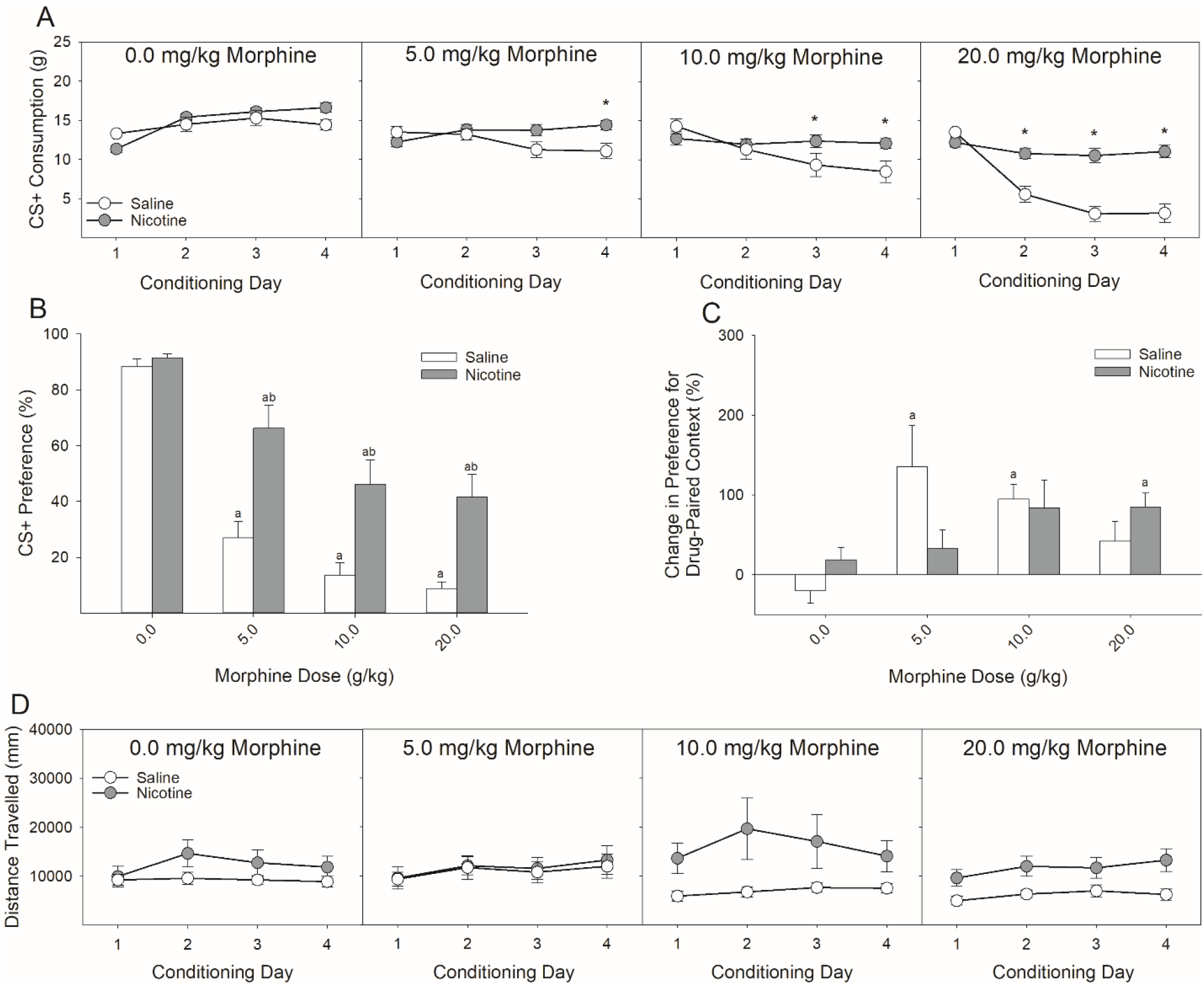
Systemic nicotine injections shift the dose-response curve of morphine-induced conditioned taste avoidance and place preference rightward. (A) Systemic pretreatment with 0.4 mg/kg of nicotine reduced the acquisition of CTA induced by three doses of morphine relative to saline pretreatment. (B) Similarly, previously nicotine-treated rats expressed a reduced CTA to saccharin following pairing with three doses of morphine. At each dose of morphine, previously nicotine-treated rats displayed significantly greater preference for the saccharin CS relative to saline-treated rats. These differences were observed in 24-h two-bottle preference tests in the absence of fluid restriction and ongoing nicotine administration. (C) Nicotine pretreatment similarly shifted the dose-response function in the CPP paradigm such that the lowest dose of morphine tested (5 mg/kg) failed to produce a significant place preference in nicotine-treated rats while sufficiently producing a place preference in saline-treated rats. At the highest dose of morphine (20 mg/kg), saline-treated rats failed to display a significant CPP, while nicotine-treated rats did. (D) There were no main or interactive effects of nicotine on the locomotion during conditioning with morphine. a’s indicate significant differences between conditioned rats and their unconditioned controls (*P*s < 0.05) and *’s and b’s indicate significant nicotine and vehicle group differences (*P*s < 0.05).

Analyses of the expression tests (Fig 3b) supported the acquisition data. A Drug x Dose ANOVA conducted on the preference for the CS+ in the absence of any further drug administrations revealed a significant 2-way interaction (*F*_(3,64)_ = 3.19, *P* < 0.05). Post-hoc analyses determined that, compared to unconditioned controls, both nicotine- and saline-treated rats demonstrated a significant decrease in the preference for saccharin at all three doses of morphine (*P*s < 0.05). Importantly, nicotine-treated rats displayed a significantly larger preference for saccharin following conditioning with all three doses of morphine relative to saline-treated rats (*Ps* < 0.05; Fig 3b). There was no effect of nicotine on the preference for saccharin in the unconditioned controls.

#### Conditioned Place Preference

Systemic nicotine administration (0.4 mg/kg) altered the pattern of expression of CPP following conditioning with multiple doses of morphine (Fig 3c). A Drug x Dose ANOVA conducted on the change in expression for preference of the drug-paired context revealed a significant 2-way interaction (*F*_(3,64)_ = 2.94, *P* < 0.05). Post-hoc analyses revealed that nicotine pretreatment shifted the dose-response curve to the right, relative to saline pretreatment. Specifically, saline-treated rats showed a significant enhancement of CPP following conditioning with 5.0, and 10 mg/kg morphine. Nicotine-treated rats failed to show a significant CPP to morphine at all doses except for the highest dose tested (20 mg/kg), a dose at which saline-treated rats did not display CPP. The CPP occurring following conditioning with 10 mg/kg in nicotine-treated rats did not survive Bonferroni correction. Peak conditioning in saline-treated rats occurred at the 5.0 mg/kg dose and peak conditioning in nicotine-treated rats occurred at the 20 m/kg dose. There was a main effect of nicotine on locomotor activity (Fig 3d) during morphine conditioning (*F*_(1,64)_ = 7.86, *P* < 0.01) but no interactive effects between nicotine and morphine.

### Experiment 3

#### Histology

Only animals for which we could determine that both cannulae were correctly placed into the insular cortex were included in statistical analyses. Fig 4a represents the placement of all injection sites across each experimental condition. Fig 4b depicts a representative image of a thionin-stained coronal section illustrating cannulae and injector track marks targeting the agranular insular cortex.

**Figure 4 -.**
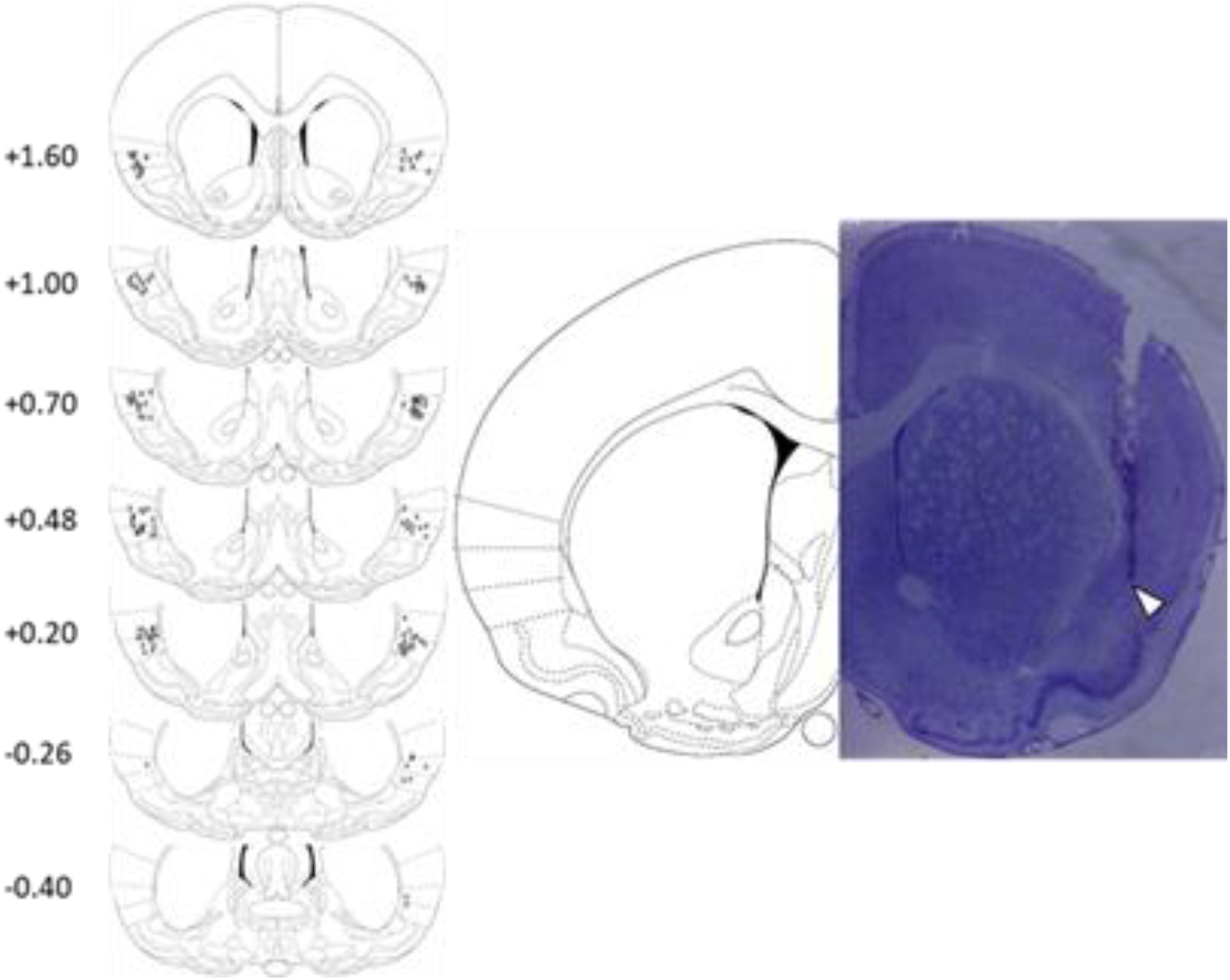
Location of injection sites within the insular cortex. A.) Coronal plates adapted from ^74^ depicting approximate injection sites from all rats included in Experiment 2. B.) Representative thionin-stained coronal section depicting the cannulae track targeting the insular cortex.

#### Conditioned Taste Avoidance

Nicotine (4 μg) infused directly into the insular cortex completely blocked the acquisition of morphine-induced CTA at all three doses of morphine (5.0, 10, & 20 mg/kg; Fig 5a & b). A Drug x Dose x Day ANOVA conducted on the intake of the CS+ during the acquisition phase of the experiment revealed a significant 3-way interaction (*F*_(9,174)_ = 3.23, *P* < 0.01). Post-hoc analyses indicated that IC-Nic, relative to IC-Veh, rats drank significantly more saccharin on days 3 and 4 of conditioning with 5.0 mg/kg morphine and on days 2, 3, and 4 of conditioning with both 10 and 20 mg/kg morphine.

**Figure 5 -.**
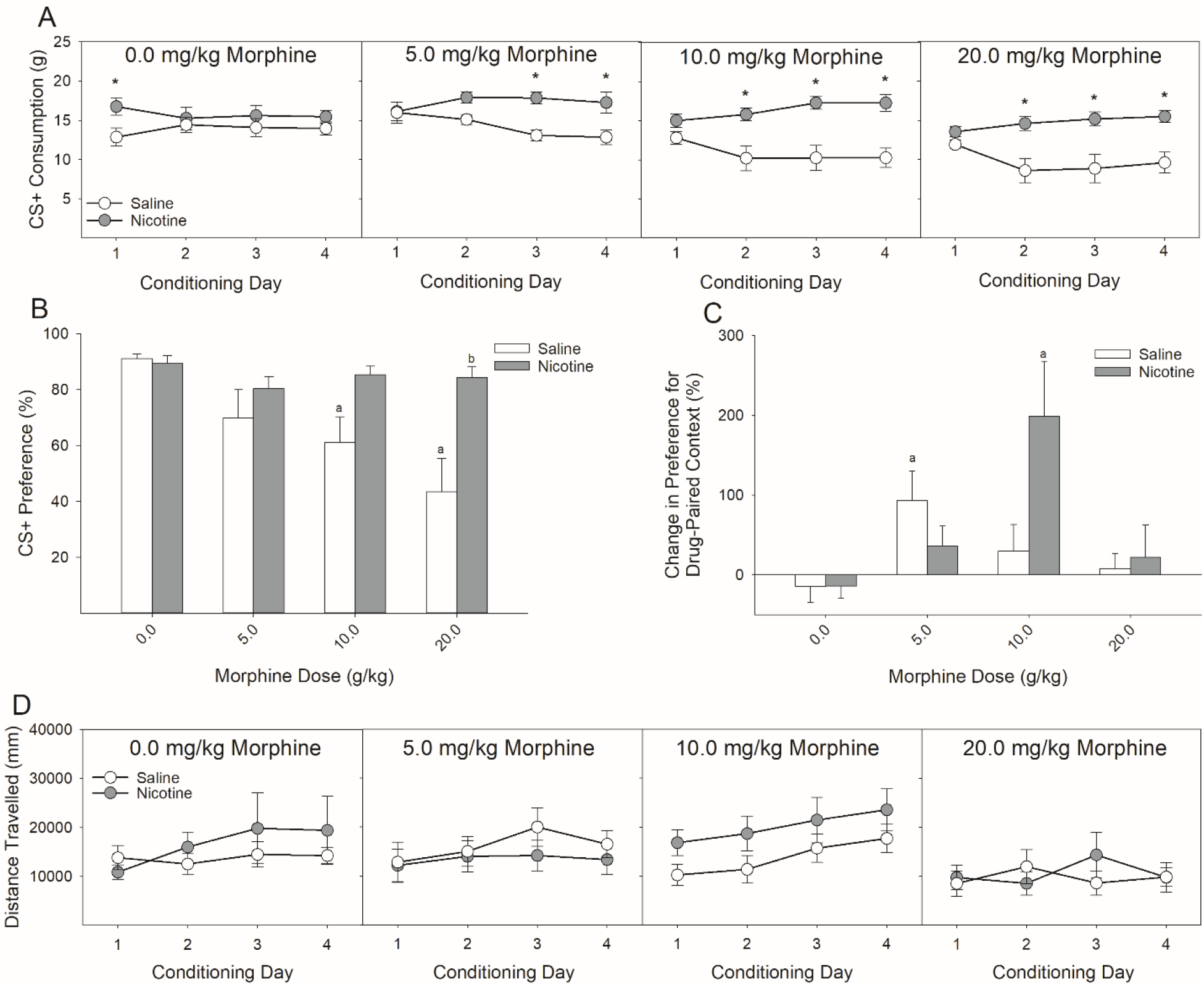
Intra-Insular nicotine infusions, similar to systemic nicotine injections, shift the dose-response curve for the acquisition of morphine-induced conditioned taste avoidance and place preference rightward. (A) Pretreatment with 4 μg of nicotine into the insular cortex blocked the acquisition of CTA induced by three doses of morphine relative to vehicle pretreatment. (B) Similarly, previously nicotine-treated rats failed to express a CTA to saccharin following pairing with any of three doses of morphine. Furthermore, at the highest dose of morphine (20 mg/kg), previously nicotine-treated rats displayed significantly greater preference for the saccharin CS relative to saline-treated rats. These differences were observed in 24-h two-bottle preference tests in the absence of fluid restriction and ongoing nicotine administration. (C) Intra-insular nicotine pretreatment similarly shifted the dose-response function in the CPP paradigm such that the lowest dose of morphine tested (5 mg/kg) failed to produce a significant place preference in nicotine-treated rats while sufficiently producing a place preference in saline-treated rats. At the 10 mg/kg dose of morphine saline-treated rats failed to display a significant CPP while nicotine-treated rats did. (D) There were no main or interactive effects of intra-insular nicotine on the locomotion during conditioning with morphine. a’s indicate significant differences between conditioned rats and their unconditioned controls (*P*s < 0.05) and *’s and b’s indicate significant nicotine and vehicle group differences (*P*s < 0.05).

Analyses of the expression test confirmed the effects observed during the acquisition phase in that IC-Nic rats failed to express a morphine-induced CTA at any dose tested, while IC-Veh rats expressed a dose-dependent decrease in the preference for saccharin over water (Fig 5b). A Drug x Dose ANOVA conducted on the preference for the CS+ in the absence of any further drug administrations revealed a significant 2-way interaction (*F*_(3,58)_ = 3.59, *P* < 0.05). Post-hoc analyses determined that, compared to unconditioned controls, IC-Veh rats expressed a CTA at all three doses of morphine while IC-Nic rats did not differ from their unconditioned controls at any dose. In addition, saline-treated rats displayed a significantly lower preference relative to nicotine-treated rats following conditioning with 20 mg/kg morphine, the difference following conditioning with 10 mg/kg morphine failed to survive Bonferroni correction. There was no effect of nicotine on the preference for saccharin in the unconditioned controls.

#### Conditioned Place Preference

Insular infusion of nicotine significantly altered the pattern of expression of CPP following conditioning with multiple doses of morphine (Fig 5c). A Drug x Dose ANOVA conducted on the change score in preference for the drug-paired floor context following conditioning with each dose of morphine revealed a significant 2-way interaction (*F*_(3,59)_ = 3.30, *P* < 0.05). Post-hoc analyses indicated that, while IC-Nic rats failed to show a reliable CPP following conditioning with 5.0 mg/kg morphine, they did show a significant CPP following conditioning with 10 mg/kg morphine, where vehicle-treated rats did not (*P* < 0.05), indicative of a rightward-ward shift in the sensitivity to morphine in the place conditioning procedures. Peak CPP in IC-Veh rats was observed following conditioning with 5.0 mg/kg morphine, relative to unconditioned controls, while peak CPP in IC-Nic rats was observed following conditioning with 10 mg/kg morphine. There were no main or interactive effects of nicotine on the locomotor response during conditioning with morphine (Fig 5d).

## Discussion

Within the present set of experiments, we demonstrate that nicotine pretreatment significantly impacts both the operant self-administration of remifentanil and morphine (Fig 1) as well as the formation of memories associated with the aversive (Fig 3a & 5a) and rewarding (Fig 3b & 5b) properties of passively-administered morphine. While previous reports have indicated that systemic nicotine can impact the formation of drug-associated memories ^12,22,31^ we have replicated those findings here and further demonstrated that nicotine, acting solely within the IC, is sufficient to produce nearly identical effects implicating the IC as an important locus of action for nicotine-induced modulation of drug-associated memories.

Systemic nicotine administration enhances the self-administration of multiple classes of commonly abused drugs in rodent models ^20,51,52^. Surprisingly, there has been remarkably little work on nicotine facilitation of opioid self-administration in preclinical models despite the abundance of evidence in humans that concurrent nicotine use is strongly associated with development of OUDs ^2,5,6^ and directly enhances consumption of methadone ^14^. Here, we found that systemic pretreatment with nicotine (0.4 mg/kg), compared to saline, markedly enhanced intake of the ultra short-acting synthetic opioid remifentanil, doubling the amount of remifentanil self-administered across a two-hour session (131.91 ± 11.24 vs 279.82 ± 28.79 μg/kg). Given the rapid metabolism of remifentanil and resultant short duration of its effects, we chose to examine the enhancement effects of nicotine on a longer lasting opioid in morphine. Similarly, nicotine pretreatment enhanced intake of morphine (4.63 ± 0.79 vs 7.13 ± 1.24 mg/kg) indicating that the enhancement by nicotine was not entirely dependent on the short duration of remifentanil. These data are consistent with findings that concurrent nicotine use increases the propensity to misuse prescribed opioid medication ^6,53^ Undoubtedly, there are likely multiple mechanisms, presumably acting in concert, through which nicotine may enhance opiate self-administration. For instance, nicotine can enhance the incentive salience of discrete drug-associated cues ^19,24,25^ and therefore increase the likelihood of drug intake upon presentation of said cues. In addition, nicotine can enhance dopamine dynamics in response to subsequently presented drugs ^54^, including disinhibition of dopamine release from the VTA ^55^, which presumably would increase the reinforcing efficacy of any subsequently administered drug. As such, nicotine pretreatment can alter the response to subsequent drugs of abuse through myriad mechanisms all independently contributing to the enhanced SUD liability observed in habitual users of nicotine-containing products ^1-5^.

One potentially novel mechanism for enhanced SUD liability resultant from concurrent nicotine use is that nicotine may promote insular dysfunction by limiting excitatory synaptic plasticity within the IC through the activation of nAchRs on GABAergic interneurons ^46-48^. Bath application of nicotine to insular slices *in vitro* facilitates GABAergic transmission and ultimately results in long-term depression (LTD) at pyramidal synapses. The IC is critically involved in the processing and integration of the interoceptive properties of drugs of abuse as well as the formation of drug-associated memories ^43-45,56-59^. Furthermore, IC dysfunction underlies many of the maladaptive decision processes that are characteristic of SUDs, including continued drug use despite adverse consequences, and is critically associated with various forms of relapse ^60-62^. In addition, the IC appears to be critical for maintenance of nicotine self-administration ^38,40,50^. Here, we demonstrate for the first time that local IC application of nicotine *in vivo* significantly impacts the formation of contextual memories associated with the aversive and reinforcing properties of opiates as would be predicted from *in vitro* slice preparations ^46^. Detrimental effects of nicotine pretreatment on contextual conditioning have been demonstrated previously, both directly and indirectly, across multiple paradigms ^11,31,63-65^. For instance, acute nicotine pretreatment in the ventral hippocampus impairs contextual fear conditioning ^64^ without affecting acute freezing signaled by a discrete cue ^65^. Additionally, adolescent exposure to nicotine impairs learning in adulthood about contexts associated with foot-shock, but has no effect on learning that a discrete cue predicts foot-shock ^63^. Adolescent exposure to nicotine similarly impairs aversive taste conditioning induced by alcohol later in adulthood ^22^. Moreover, the context associated with nicotine self-administration is not sufficient to maintain high-levels of nicotine self-administration in the absence of discrete cues suggesting that nicotine administration may particularly promote the salience of discrete cues at the expense of contextual information ^66^.

In the present study we examined the effects of both systemic (0.4 mg/kg) and intra-insular (4 μg) pretreatment with nicotine on a combined CTA/CPP paradigm in which we measured the strength of both place and taste conditioning in the same animal following four passive administrations of morphine. We tested multiple doses of morphine (5.0, 10, & 20 mg/kg) across individual groups of rats and found that administration of nicotine, relative to vehicle, interfered with the acquisition of CTA to all doses of morphine tested. This response was observed during the acquisition phase of the experiment when the animals were water deprived (Fig 3a & 5a) and actively receiving nicotine infusions, and also in the expression phase in which the animals had 24-h access to fluid and were no longer receiving nicotine administrations (Fig 3b & 5b). These results indicate that the effects were not due to any ability for nicotine to induce hyperphagia of the saccharin solution or that nicotine was somehow interacting with deprivation state to influence greater fluid consumption. Similarly, we found that that both forms of nicotine administration impacted morphine CPP in manner that indicated a similar rightward shift in the dose-response curve. Consistent with previous studies ^11,30^, we show that systemic nicotine administration blocked the CPP observed following conditioning with a low dose of morphine (5.0 mg/kg), and this was also observed following intra-insular nicotine pretreatment. Higher doses of morphine that failed to condition a significant CPP in saline-treated rats, produced a significant CPP following both systemic and intra-insular nicotine administration (20 and 10 mg/kg, respectively).

This pattern of results was remarkably similar across both nicotine administration regimens. Specifically, nicotine treatment shifted the dose response curve for morphine in both CTA and CPP paradigms rightward indicative of decreased sensitivity to the effects of morphine in these specific paradigms. Consistent with our previous findings ^12^, nicotine did not result in an inability to form these associations, but rather interfered with the strength of conditioning, and these effects closely mimic those observed following IC lesions and pharmacological inactivation ^43,45^ as well as intracerebroventricular nicotine administration ^30^. In agreement with these previous studies, comparisons of the data from experiments 2 and 3 suggest that the animals with insular cannulae in experiment 3, regardless of pretreatment condition, appeared to be less sensitive to the properties of morphine in the contextual conditioning paradigms relative to the intact animals of experiment 2. Given that implantation of the bilateral guide cannulae and infusions undoubtedly produced some damage to insular tissue, this observation is consistent with previous lesion studies. Regardless, across both experiments, nicotine produced similar effects following both routes of administration, compared to vehicle administration. Given that systemic nicotine would invariably be acting across numerous brain areas, the finding that the pattern of results between systemic and intra-insular administration were largely similar suggests that IC is an important area for the effects of nicotine on drug-associated contextual conditioning. It would be important to determine in future studies the relative involvement of additional brain areas associated with these forms of conditioning (e.g. VTA, hippocampus, amygdala etc. ^67^) as well as blocking insular nAChRs following systemic nicotine administration to determine the necessity IC nAChRs to the present findings.

The present pattern of results reveals two important considerations. The first is that CTA and CPP conditioning are, in fact, dissociable. For instance, we found that IC nicotine administration blocked CTA to a given dose of morphine (10 mg/kg) while simultaneously enhancing CPP to that same dose. Given our findings and those of others ^68^, it follows that CTA and CPP are conditioned by differing interoceptive properties of morphine. The second consideration is that nicotine-treated animals were not incapable of forming opiate-associated memories; rather they required higher doses of morphine to do so. This is consistent with our previous work on systemic nicotine and morphine-CTA ^12^ wherein systemically-treated nicotine rats could learn to avoid a taste stimulus paired with morphine administration but they needed a substantially larger dose in order to do so comparably to saline-treated rats. Importantly, nicotine administration does not completely abolish all forms of insular-dependent associative learning; nicotine-treated rats competently learn a taste aversion to the visceral malaise-inducing compound lithium chloride ^21^ which has no known reinforcing value. This raises the intriguing possibility that through putatively enhancing insular inactivation, nicotine may be reducing the perception of the interoceptive properties of subsequent drug administration in a manner that facilitates larger, and potentially more frequent, doses. In agreement, human smokers are less sensitive to the analgesic properties of opiates ^53^ requiring higher and more frequent doses. In addition, nicotine impacts interoceptive detection of cocaine in humans such that pretreatment with nicotine necessitates longer periods and stronger doses in order to report the passive administration of cocaine ^69^, relative to placebo. In paradigms examining the interoceptive impact of combined nicotine and alcohol administration, nicotine is found to be the more salient interoceptive cue ^70^. Furthermore, insular manipulations in rats interfere with processing the interoceptive effects of alcohol ^71,72^. While our findings are not inconsistent with such an interpretation, the present studies were primarily concerned with the effects of nicotine on opioid self-administration and the formation of drug-associated memories and therefore more work is required to determine if insular nicotinic activity results in acute decrements in drug detection and these studies are currently being conducted.

These current studies are not without their limitations. For instance, all of these studies were conducted in male rats. In a previous report ^12^, we demonstrated that systemic nicotine impaired the acquisition of ethanol CTA in both males and females, and the effect appeared to be larger in female rats. It remains to be determined if the IC similarly contributes to this effect in females. Additionally, while these forms of site-specific behavioral pharmacological approaches can yield insight into the relative involvement of a brain area towards a phenomenon, they are limited in their specificity of various sub-regions of the structure in question. Specifically, it is certain that in this, and other experimental approaches, the drug infusions spread throughout the brain area and so we cannot make claims as to the contribution of insular subregions (i.e. granular, dysgranular, or agranular) to the present results. As such, future studies using these behavioral paradigms and employing circuit specific viral approaches ^73^ are warranted. Finally, though the combined CTA/CPP approach is powerful in determining the strength of approach and avoidance inclinations in a given animal at a given dose of drug, there is some room for interactive effects between the two conditioning paradigms. For instance, it is possible the availability of saccharin could serve as an occasion setter and thus influence the strength of subsequent conditioning in the CPP context. It would be informative to determine if the present results hold if both paradigms are used independently. In support of such an assumption, we have previously demonstrated that systemic nicotine pretreatment similarly impairs CTA acquisition in the absence of CPP conditioning ^21^ and others have shown identical effects of systemic and intracerebroventricular nicotine in CPP paradigms, at least following conditioning with 5.0 mg/kg morphine, in the absence of antecedent saccharin CTA ^11,30^.

In summary, we report that acute nicotine administration significantly enhances the self-administration of remifentanil and morphine. In addition, acute nicotine administered either systemically or directly to the IC modulates the formation of drug-associated memories in a manner that indicates that nicotine-treated rats require higher doses in order to form comparable contextual drug associations relative to controls. These data not only emphasize the importance of nicotine co-use in the development and maintenance of problematic drug use but also further our understanding of the importance of the insular cortex in nicotine-facilitated polysubstance use.

## Methods and Materials

### Animals and housing

161 adult male Long-Evans rats weighing approximately 350 g (Envigo; Indianapolis, IN) were individually housed on a reverse light cycle in polycarbonate cages in a humidity- and temperature-controlled vivarium. Following arrival to the facilities, rats were handled daily for three days and given at least one week to acclimate prior to the start of the experiments. 2 rats were excluded from experiment 2 due to development of illness and one rat died before completion of the study. 11 rats were eventually excluded from Experiment 2 due to either misplaced cannulae or the development of illness. All procedures were approved by the University at Buffalo Institutional Animal Care and Use Committee.

### Chemical stimuli

Nicotine hydrogen tartrate salt (Glentham Life Sciences; U.K.) was dissolved in phosphate buffered saline (PBS; pH 7.2-7.4). All nicotine doses are expressed as the freebase. Morphine sulfate (Spectrum Chemicals; New Brunswick, NJ) was dissolved in saline. Remifentanil hydrochloride (NIDA drug supply program) was dissolved in saline. Saccharin sodium (Sigma-Aldrich; St Louis, MO) was dissolved in tap water.

## Experiment 1: Does systemic nicotine administration enhance intravenous morphine self-administration?

### Surgical procedures

Rats (n=9) were surgically implanted with chronically indwelling catheters in the right jugular vein under isoflurane anesthesia (1-3%). Briefly, a 15cm catheter line (C30PU-RJV1402, Instech) was implanted into the right external jugular vein and the other end was connected to an externalized vascular access button (VABR1B/22, Instech). Each access button was implanted just posterior to the scapular region on the dorsal side of the rat. Catheters were flushed with 0.1 mL of heparinized saline and enrofloxacin (Baytril, Bayer HealthCare LLC; Shawnee Mission, KS) before and after self-administration on test days. Catheter patency was verified at the conclusion of the study by flushing 0.1 ml of ketamine (10 mg/ml) and verifying the expression of ataxia.

### Behavioral procedures: Intravenous self-administration

Rats were trained to press a lever in standard operant boxes (Med Associates; St Albans, VT) for delivery of a drug reinforcer in two-hour sessions. Behavior was shaped such that one press of the active-lever resulted in intravenous delivery of remifentanil (3.2 μg/kg) while presses of the inactive-lever had no consequence. Once stable responding was established (7 days), rats were advanced to an FR-3 schedule of reinforcement. Once stable responding was confirmed on FR-3, rats were given two, non-contingent sub cutaneous (s.c.) injections of nicotine (0.4 mg/kg) across two days in their homecage, in the absence of operant test sessions, in order to familiarize them to the effects of nicotine. Next, rats were tested for FR-3 responding for remifentanil in two sessions, once each under the effects of nicotine pretreatment (0.4 mg/kg s.c., 15 min before session start) or saline (1.0 ml/kg s.c., 15 min before session start). One week following the last FR-3 session with remifentanil, rats were placed back into the operant chambers and now allowed to self-administer morphine (0.3 mg/kg) on an FR-3 schedule. Once stable responding for morphine was confirmed the rats were similarly tested for FR-3 morphine responding following nicotine (0.4 mg/kg) and saline pretreatment. Location of the active and inactive lever and order of testing (nicotine or saline) were counter-balanced across all rats.

## Experiment 2: Does systemic nicotine alter contextual conditioning with the interoceptive effects of morphine?

### Behavioral procedures: Combined CTA/CPP paradigm

Rats (n=72) were maintained on a 23 h water deprivation schedule and tested in a combined CTA/CPP paradigm (see timeline; Fig 2a). The behavioral paradigm consisted of 4 distinct phases. In order of presentation, they were: drug pretreatment (nicotine/saline); fluid access (saccharin/water); US administration (0.0, 5.0, 10, 20 mg/kg morphine); place conditioning. Briefly, rats were trained to consume fluid during a 30 min session until intakes stabilized (~5-7 days). On the last training day, immediately following the fluid-access period, rats were placed in the CPP apparatus and their baseline preference for the two contexts was assessed. We employed a biased CPP procedure where the least preferred floor from the baseline assessment was assigned as the CS+ condition and the most preferred floor as the CS-condition. Immediately following training, rats were assigned to an experimental condition in a counter-balanced fashion.

**Figure 2 -.**
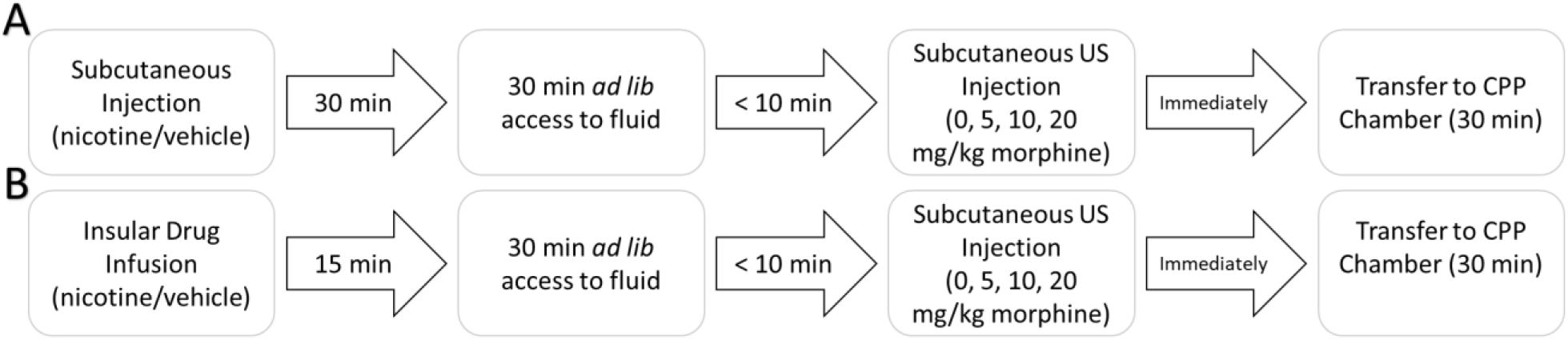
Experimental timeline on conditioning days. On conditioning days in Experiment 2 (A), rats were first subcutaneously injected with their assigned drug and returned to their home cage. 15 min later the rats were given access to fluid in their homecage for 30 min. Within 10 minutes of cessation of the fluid access period, rats were injected with their assigned morphine dose (*s.c.*) and immediately placed into the CPP chambers for 30 min prior to being returned to their homecage. In Experiment 3 (B) the same protocol was followed with the exception that the animals were bilaterally infused into the insula with their assigned drug and returned to their home cage. 15 min later the rats were given access to fluid in their homecage and all other procedures were identically followed.

During testing, rats were first injected subcutaneously with either 0.4 mg/kg nicotine or saline. Following injections, rats were returned to their homecage for 30 min. Next. They were given access to fluid (saccharin or water) for 30 min in their homecage. Following the fluid-access period, rats were removed from their homecage and injected with either saline or morphine (5.0, 10, or 20 mg/kg, s.c.) and placed into the CPP apparatus equipped with the appropriate floor context for 30 min. This resulted in a total of 8 experimental groups: three doses of morphine following systemic nicotine and the same three doses following saline and their respective unconditioned controls.

On CS-days, all rats were given access to water for 30 min immediately followed by an injection of saline prior to transfer to the CPP chambers. On CS+ days, all rats were given access to 0.1 % saccharin for 30 min and immediately injected with their assigned morphine dose prior to transfer to the CPP chambers (see Fig 2a). Conditioning lasted for a total of 8 days in which CS- and CS+ conditions were alternated for a total of four taste-context-drug pairings.

Following conditioning, the strength of the place preference was first assessed by placing the rats in the CPP apparatus equipped with both floor contexts in the absence of any drug administrations. Rats were left for 30 min and the amount of time spent in either context was measured. One day after CPP expression tests, the strength of CTA was assessed by giving rats 24 h access to two fluid bottles in their homecage, one bottle contained tap water and the other contained 0.1 % saccharin. Rats were free to consume from either bottle and the preference for saccharin over water was calculated.

## Experiment 3: Does nicotine delivered locally to the insular cortex affect contextual interoceptive conditioning with morphine?

### Surgical procedures

Rats (n=78) were restrained in a stereotaxic apparatus (Kopf Instruments; Tujunga, CA) with non-traumatic ear bars and maintained on isoflurane (2-4%) throughout the duration of the surgical procedure. Stainless-steel guide cannulae (26 gauge, 8.0 mm in length; Invivo1; Roanoke, VA) targeting 2.0 mm dorsal to the agranular insular cortex (+0.5 A/P; ±5.7 M/L; -4.4 D/V) were bilaterally implanted and fixed to the skull with screws and dental cement. Dummy cannulae (36 gauge) were inserted into the guide cannulae in order to prevent occlusion. All rats were given 7 days to recover prior to the start of behavioral experiments.

Following recovery, rats were trained and tested in an identical combined CTA/CPP design as described above (Experiment 2) with a few critical changes (Figure 2b). Each day, rats were gently restrained for approximately two minutes while dummy cannulae were removed and injectors were inserted into both cannulae. Immediately following training, rats were assigned to an experimental condition in a counter-balanced fashion. During testing, rats were first bilaterally infused into the IC with 1 μl of their assigned stimulus (4 μg nicotine; IC-Nic or vehicle; IC-Veh) at a rate of 0.5 μl/min. Injectors (extending 2.0 mm beyond cannulae) were left in place for an additional minute to allow for adequate diffusion away from the injection tract. Following infusions, rats were returned to their homecage for 15 min before given access to fluid (saccharin or water) for 30 min. Following the fluid-access period, rats were removed from their home cage and injected with either saline or morphine (5.0, 10, or 20 mg/kg, s.c.) and placed into the CPP apparatus equipped with the appropriate floor context for 30 min. This resulted in a total of 8 experimental groups: three doses of morphine following IC-Veh and three doses following IC-Nic and their respective unconditioned controls. Testing lasted for 8 days (4 alternations of CS- and CS+ conditioning trials) and the strength of CTA and CPP conditioning as a function of insular-drug delivery and morphine dose (0.0, 5.0, 10, or 20 mg/kg, s.c.) were assessed as described above.

### Histology

Following behavioral testing, rats were anesthetized with pentobarbital and transcardially perfused with ice cold saline followed by 4% paraformaldehyde (PFA) in 0.1 M phosphate buffer and their brains were removed. Brains were post-fixed in PFA overnight. Fixed brains were cryoprotected by sinking in a 15% w/v buffered sucrose solution for one day followed by a 30% sucrose solution. The insular cortex was sectioned at 40 μm on a freezing microtome (Hacker Instruments; Winnsboro, SC) and sections were transferred to slides and stained with a 0.1% thionin solution. Images were obtained of the brain sections on a microscope equipped with a digital camera (Cole-Parmer; Vernon Hills, IL) and placement of injection sites within the insular cortex were verified (see Fig 4).

## Data Analyses

For the IVSA portion of the experiments, the total number of active and inactive lever presses, as well as total infusions earned were independently analyzed for both remifentanil and morphine IVSA in repeated measures ANOVAs with Drug pretreatment (nicotine or saline) as within-subjects factor. Post-hoc analyses of repeated factors were conducted with Fisher’s LSD.

For the CTA portion of the experiments, intake of the saccharin CS and water were analyzed independently in a mixed repeated measures ANOVA with Drug (nicotine or saline) and morphine Dose (0.0, 5.0, 10, 20 mg/kg morphine) as between-subjects factors and conditioning Day as within-subjects factors. Post-hoc analyses of repeated factors was conducted with Fisher’s LSD. For CTA 2-bottle expression tests, intake data were converted to a preference score (CS+ over CS-) and were analyzed with a two-factor (Drug and morphine Dose) between-subjects ANOVA. Bonferroni-corrected t-tests were used to assess strength of conditioning within- and between-groups separately.

For CPP expression tests, the percent change in preference for the drug-paired context relative to the baseline preference test was analyzed with a two-factor (Drug and US Dose) between-subjects ANOVA. Separate Bonferroni-corrected t-tests were used to assess strength of conditioning within- and between-groups. Statistical analyses were conducted in Statistica 12 software package.

## Author contributions

GCL and PJM designed experiments, GCL and CPK conducted experiments and analyzed data; GCL, CPK, and PJM wrote the manuscript.

## References

1 Weinberger, A. H. et al. Cigarette Smoking Is Associated With Increased Risk of Substance Use Disorder Relapse: A Nationally Representative, Prospective Longitudinal Investigation. J Clin Psychiatry 78, e152–e160, doi:10.4088/JCP.15m10062 (2017).

2 Nahvi, S., Richter, K., Li, X., Modali, L. & Arnsten, J. Cigarette smoking and interest in quitting in methadone maintenance patients. Addict Behav 31, 2127–2134, doi:10.1016/j.addbeh.2006.01.006 (2006).

3 Kohut, S. J. Interactions between nicotine and drugs of abuse: a review of preclinical findings. Am J Drug Alcohol Abuse 43, 155–170, doi:10.1080/00952990.2016.1209513 (2017).

4 Tuesta, L. M., Fowler, C. D. & Kenny, P. J. Recent advances in understanding nicotinic receptor signaling mechanisms that regulate drug self-administration behavior. Biochemical pharmacology 82, 984–995, doi:10.1016/j.bcp.2011.06.026 (2011).

5 Frosch, D. L., Shoptaw, S., Nahom, D. & Jarvik, M. E. Associations between tobacco smoking and illicit drug use among methadone-maintained opiate-dependent individuals. Exp Clin Psychopharm 8, 97–103, doi:Doi 10.1037/1064-1297.8.1.97 (2000).

6 Zale, E. L. et al. Tobacco Smoking, Nicotine Dependence, and Patterns of Prescription Opioid Misuse: Results From a Nationally Representative Sample. Nicotine Tob Res 17, 1096–1103, doi:10.1093/ntr/ntu227 (2015).

7 Yoon, J. H., Lane, S. D. & Weaver, M. F. Opioid Analgesics and Nicotine: More Than Blowing Smoke. J Pain Palliat Care Pharmacother 29, 281–289, doi:10.3109/15360288.2015.1063559 (2015).

8 Skurtveit, S., Furu, K., Selmer, R., Handal, M. & Tverdal, A. Nicotine dependence predicts repeated use of prescribed opioids. Prospective population-based cohort study. Ann Epidemiol 20, 890–897, doi:10.1016/j.annepidem.2010.03.010 (2010).

9 Henningfield, J. E., Clayton, R. & Pollin, W. Involvement of Tobacco in Alcoholism and Illicit Drug-Use. Brit J Addict 85, 279–292 (1990).

10 Feng, B. et al. Blocking alpha4beta2 and alpha7 nicotinic acetylcholine receptors inhibits the reinstatement of morphine-induced CPP by drug priming in mice. Behav Brain Res 220, 100–105, doi:10.1016/j.bbr.2011.01.040 (2011).

11 Zarrindast, M. R., Faraji, N., Rostami, P., Sahraei, H. & Ghoshouni, H. Cross-tolerance between morphine- and nicotine-induced conditioned place preference in mice. Pharmacology, biochemistry, and behavior 74, 363–369 (2003).

12 Loney, G. C. & Meyer, P. J. Nicotine pre-treatment reduces sensitivity to the interoceptive stimulus effects of commonly abused drugs as assessed with taste conditioning paradigms. Drug Alcohol Depend 194, 341–350, doi:10.1016/j.drugalcdep.2018.07.048 (2019).

13 Glick, S. D., Ramirez, R. L., Livi, J. M. & Maisonneuve, I. M. 18-Methoxycoronaridine acts in the medial habenula and/or interpeduncular nucleus to decrease morphine self-administration in rats. European journal of pharmacology 537, 94–98, doi:10.1016/j.ejphar.2006.03.045 (2006).

14 Spiga, R., Schmitz, J. & Day, J., 2nd. Effects of nicotine on methadone self-administration in humans. Drug Alcohol Depend 50, 157–165 (1998).

15 Crombag, H. S., Bossert, J. M., Koya, E. & Shaham, Y. Review. Context-induced relapse to drug seeking: a review. Philos Trans R Soc Lond B Biol Sci 363, 3233–3243, doi:10.1098/rstb.2008.0090 (2008).

16 McHugh, R. K. et al. Assessing craving and its relationship to subsequent prescription opioid use among treatment-seeking prescription opioid dependent patients. Drug Alcohol Depend 145, 121–126, doi:10.1016/j.drugalcdep.2014.10.002 (2014).

17 Krasnova, I. N. et al. Incubation of methamphetamine and palatable food craving after punishment-induced abstinence. Neuropsychopharmacology : official publication of the American College of Neuropsychopharmacology 39, 2008–2016, doi:10.1038/npp.2014.50 (2014).

18 Venniro, M., Caprioli, D. & Shaham, Y. Animal models of drug relapse and craving: From drug priming-induced reinstatement to incubation of craving after voluntary abstinence. Prog Brain Res 224, 25–52, doi:10.1016/bs.pbr.2015.08.004 (2016).

19 Loney, G. C., Angelyn, H., Cleary, L. M. & Meyer, P. J. Nicotine Produces a High-Approach, Low-Avoidance Phenotype in Response to Alcohol-Associated Cues in Male Rats. Alcoholism, clinical and experimental research 43, 1284–1295, doi:10.1111/acer.14043 (2019).

20 Bechtholt, A. J. & Mark, G. P. Enhancement of cocaine-seeking behavior by repeated nicotine exposure in rats. Psychopharmacology 162, 178–185, doi:10.1007/s00213-002-1079-1 (2002).

21 Loney, G. C. & Meyer, P. J. Nicotine reduces the sensitivity to the interoceptive properties of commonly abused drugs as assessed with taste conditioning paradigms. Drug Alcohol Depend In Press (2018).

22 Rinker, J. A. et al. Exposure to nicotine during periadolescence or early adulthood alters aversive and physiological effects induced by ethanol. Pharmacology, biochemistry, and behavior 99, 7–16, doi:10.1016/j.pbb.2011.03.009 (2011).

23 Kunin, D., Smith, B. R. & Amit, Z. Nicotine and ethanol interaction on conditioned taste aversions induced by both drugs. Pharmacology, biochemistry, and behavior 62, 215–221 (1999).

24 Maddux, J. N. & Chaudhri, N. Nicotine-induced enhancement of Pavlovian alcohol-seeking behavior in rats. Psychopharmacology 234, 727–738, doi:10.1007/s00213-016-4508-2 (2017).

25 Stringfield, S. J., Boettiger, C. A. & Robinson, D. L. Nicotine-enhanced Pavlovian conditioned approach is resistant to omission of expected outcome. Behav Brain Res 343, 16–20, doi:10.1016/j.bbr.2018.01.023 (2018).

26 Bevins, R. A. & Palmatier, M. I. Extending the role of associative learning processes in nicotine addiction. Behav Cogn Neurosci Rev 3, 143–158, doi:10.1177/1534582304272005 (2004).

27 Palmatier, M. I., Kellicut, M. R., Brianna Sheppard, A., Brown, R. W. & Robinson, D. L. The incentive amplifying effects of nicotine are reduced by selective and non-selective dopamine antagonists in rats. Pharmacology, biochemistry, and behavior 126, 50–62, doi:10.1016/j.pbb.2014.08.012 (2014).

28 Nunes, E. J. et al. Cholinergic Receptor Blockade in the VTA Attenuates Cue-Induced Cocaine-Seeking and Reverses the Anxiogenic Effects of Forced Abstinence. Neuroscience 413, 252–263, doi:10.1016/j.neuroscience.2019.06.028 (2019).

29 Singh, P. K. & Lutfy, K. Nicotine pretreatment reduced cocaine-induced CPP and its reinstatement in a sex- and dose-related manner in adult C57BL/6J mice. Pharmacology, biochemistry, and behavior 159, 84–89, doi:10.1016/j.pbb.2017.07.010 (2017).

30 Shams, J. et al. Effects of ultra-low doses of nicotine on the expression of morphine-induced conditioned place preference in mice. Behavioural pharmacology 17, 629–635, doi:10.1097/FBP.0b013e3280102d68 (2006).

31 Araki, H. et al. Nicotine attenuates place aversion induced by naloxone in single-dose, morphine-treated rats. Psychopharmacology 171, 398–404, doi:10.1007/s00213-003-1595-7 (2004).

32 Ishida, S. et al. alpha7 Nicotinic acetylcholine receptors in the central amygdaloid nucleus alter naloxone-induced withdrawal following a single exposure to morphine. Psychopharmacology 214, 923–931, doi:10.1007/s00213-010-2101-7 (2011).

33 Ackroff, K. & Sclafani, A. Flavor preferences conditioned by intragastric ethanol with limited access training. Pharmacology, biochemistry, and behavior 75, 223–233 (2003).

34 Cunningham, C. L. & Niehus, J. S. Flavor preference conditioning by oral self-administration of ethanol. Psychopharmacology 134, 293–302, doi:10.1007/s002130050452 (1997).

35 Loney, G. C. & Meyer, P. J. Brief Exposures to the Taste of Ethanol (EtOH) and Quinine Promote Subsequent Acceptance of EtOH in a Paradigm that Minimizes Postingestive Consequences. Alcoholism, clinical and experimental research 42, 589–602, doi:10.1111/acer.13581 (2018).

36 Pitchers, K. K., Kane, L. F., Kim, Y., Robinson, T. E. & Sarter, M. ‘Hot’ vs. ‘cold’ behavioural-cognitive styles: motivational-dopaminergic vs. cognitive-cholinergic processing of a Pavlovian cocaine cue in sign- and goal-tracking rats. Eur J Neurosci 46, 2768–2781, doi:10.1111/ejn.13741 (2017).

37 Gogolla, N. The insular cortex. Curr Biol 27, R580–R586, doi:10.1016/j.cub.2017.05.010 (2017).

38 Naqvi, N. H., Rudrauf, D., Damasio, H. & Bechara, A. Damage to the insula disrupts addiction to cigarette smoking. Science 315, 531–534, doi:10.1126/science.1135926 (2007).

39 Naqvi, N. H., Gaznick, N., Tranel, D. & Bechara, A. The insula: a critical neural substrate for craving and drug seeking under conflict and risk. Annals of the New York Academy of Sciences 1316, 53–70, doi:10.1111/nyas.12415 (2014).

40 Forget, B., Pushparaj, A. & Le Foll, B. Granular insular cortex inactivation as a novel therapeutic strategy for nicotine addiction. Biol Psychiatry 68, 265–271, doi:10.1016/j.biopsych.2010.01.029 (2010).

41 Pushparaj, A. & Le Foll, B. Involvement of the caudal granular insular cortex in alcohol self-administration in rats. Behav Brain Res 293, 203–207, doi:10.1016/j.bbr.2015.07.044 (2015).

42 Jaramillo, A. A., Van Voorhies, K., Randall, P. A. & Besheer, J. Silencing the insular-striatal circuit decreases alcohol self-administration and increases sensitivity to alcohol. Behav Brain Res 348, 74–81, doi:10.1016/j.bbr.2018.04.007 (2018).

43 Geddes, R. I., Han, L., Baldwin, A. E., Norgren, R. & Grigson, P. S. Gustatory insular cortex lesions disrupt drug-induced, but not lithium chloride-induced, suppression of conditioned stimulus intake. Behavioral neuroscience 122, 1038–1050, doi:10.1037/a0012748 (2008).

44 Contreras, M., Ceric, F. & Torrealba, F. Inactivation of the interoceptive insula disrupts drug craving and malaise induced by lithium. Science 318, 655–658, doi:10.1126/science.1145590 (2007).

45 Li, C. L., Zhu, N., Meng, X. L., Li, Y. H. & Sui, N. Effects of inactivating the agranular or granular insular cortex on the acquisition of the morphine-induced conditioned place preference and naloxone-precipitated conditioned place aversion in rats. J Psychopharmacol 27, 837–844, doi:10.1177/0269881113492028 (2013).

46 Sato, H., Kawano, T., Yin, D. X., Kato, T. & Toyoda, H. Nicotinic activity depresses synaptic potentiation in layer V pyramidal neurons of mouse insular cortex. Neuroscience 358, 13–27, doi:10.1016/j.neuroscience.2017.06.031 (2017).

47 Toyoda, H. Role of nicotinic acetylcholine receptors for modulation of microcircuits in the agranular insular cortex. J Oral Biosci 61, 5–11, doi:10.1016/j.job.2018.12.001 (2019).

48 Toyoda, H. Nicotine facilitates synaptic depression in layer V pyramidal neurons of the mouse insular cortex. Neurosci Lett 672, 78–83, doi:10.1016/j.neulet.2018.02.046 (2018).

49 Jiang, C. et al. Morphine coordinates SST and PV interneurons in the prelimbic cortex to disinhibit pyramidal neurons and enhance reward. Mol Psychiatry, doi:10.1038/s41380-019-0480-7 (2019).

50 Hollander, J. A., Lu, Q., Cameron, M. D., Kamenecka, T. M. & Kenny, P. J. Insular hypocretin transmission regulates nicotine reward. Proceedings of the National Academy of Sciences of the United States of America 105, 19480–19485, doi:10.1073/pnas.0808023105 (2008).

51 Cortright, J. J., Sampedro, G. R., Neugebauer, N. M. & Vezina, P. Previous exposure to nicotine enhances the incentive motivational effects of amphetamine via nicotine-associated contextual stimuli. Neuropsychopharmacology : official publication of the American College of Neuropsychopharmacology 37, 2277–2284, doi:10.1038/npp.2012.80 (2012).

52 Le, A. D., Wang, A., Harding, S., Juzytsch, W. & Shaham, Y. Nicotine increases alcohol self-administration and reinstates alcohol seeking in rats. Psychopharmacology 168, 216–221, doi:10.1007/s00213-002-1330-9 (2003).

53 Steinmiller, C. L. et al. Postsurgical patient-controlled opioid self-administration is greater in hospitalized abstinent smokers than nonsmokers. J Opioid Manag 8, 227–235, doi:10.5055/jom.2012.0120 (2012).

54 Zernig, G., O’Laughlin, I. A. & Fibiger, H. C. Nicotine and heroin augment cocaine-induced dopamine overflow in nucleus accumbens. European journal of pharmacology 337, 1–10 (1997).

55 Mansvelder, H. D., Keath, J. R. & McGehee, D. S. Synaptic mechanisms underlie nicotine-induced excitability of brain reward areas. Neuron 33, 905–919, doi:10.1016/s0896-6273(02)00625-6 (2002).

56 Naqvi, N. H. & Bechara, A. The insula and drug addiction: an interoceptive view of pleasure, urges, and decision-making. Brain Struct Funct 214, 435–450, doi:10.1007/s00429-010-0268-7 (2010).

57 Goldstein, R. Z. et al. The neurocircuitry of impaired insight in drug addiction. Trends Cogn Sci 13, 372–380, doi:10.1016/j.tics.2009.06.004 (2009).

58 Moschak, T. M., Wang, X. & Carelli, R. M. A Neuronal Ensemble in the Rostral Agranular Insula Tracks Cocaine-Induced Devaluation of Natural Reward and Predicts Cocaine Seeking. The Journal of neuroscience : the official journal of the Society for Neuroscience 38, 8463–8472, doi:10.1523/JNEUROSCI.1195-18.2018 (2018).

59 Zito, K. A., Bechara, A., Greenwood, C. & van der Kooy, D. The dopamine innervation of the visceral cortex mediates the aversive effects of opiates. Pharmacology, biochemistry, and behavior 30, 693–699 (1988).

60 Paulus, M. P., Rogalsky, C., Simmons, A., Feinstein, J. S. & Stein, M. B. Increased activation in the right insula during risk-taking decision making is related to harm avoidance and neuroticism. Neuroimage 19, 1439–1448 (2003).

61 Stewart, J. L. et al. You are the danger: attenuated insula response in methamphetamine users during aversive interoceptive decision-making. Drug Alcohol Depend 142, 110–119, doi:10.1016/j.drugalcdep.2014.06.003 (2014).

62 Stewart, J. L., Butt, M., May, A. C., Tapert, S. F. & Paulus, M. P. Insular and cingulate attenuation during decision making is associated with future transition to stimulant use disorder. Addiction 112, 1567–1577, doi:10.1111/add.13839 (2017).

63 Spaeth, A. M., Barnet, R. C., Hunt, P. S. & Burk, J. A. Adolescent nicotine exposure disrupts context conditioning in adulthood in rats. Pharmacology, biochemistry, and behavior 96, 501–506, doi:10.1016/j.pbb.2010.07.011 (2010).

64 Kenney, J. W., Raybuck, J. D. & Gould, T. J. Nicotinic receptors in the dorsal and ventral hippocampus differentially modulate contextual fear conditioning. Hippocampus 22, 1681–1690, doi:10.1002/hipo.22003 (2012).

65 Kutlu, M. G., Oliver, C. & Gould, T. J. The effects of acute nicotine on contextual safety discrimination. J Psychopharmacol 28, 1064–1070, doi:10.1177/0269881114552743 (2014).

66 Caggiula, A. R. et al. Cue dependency of nicotine self-administration and smoking. Pharmacology, biochemistry, and behavior 70, 515–530 (2001).

67 Subramaniyan, M. & Dani, J. A. Dopaminergic and cholinergic learning mechanisms in nicotine addiction. Ann N Y Acad Sci 1349, 46–63, doi:10.1111/nyas.12871 (2015).

68 Verendeev, A. & Riley, A. L. Relationship between the rewarding and aversive effects of morphine and amphetamine in individual subjects. Learn Behav 39, 399–408, doi:10.3758/s13420-011-0035-5 (2011).

69 Kouri, E. M., Stull, M. & Lukas, S. E. Nicotine alters some of cocaine’s subjective effects in the absence of physiological or pharmacokinetic changes. Pharmacology, biochemistry, and behavior 69, 209–217 (2001).

70 Randall, P. A., Cannady, R. & Besheer, J. The nicotine + alcohol interoceptive drug state: contribution of the components and effects of varenicline in rats. Psychopharmacology 233, 3061–3074, doi:10.1007/s00213-016-4354-2 (2016).

71 Jaramillo, A. A., Randall, P. A., Frisbee, S. & Besheer, J. Modulation of sensitivity to alcohol by cortical and thalamic brain regions. Eur J Neurosci 44, 2569–2580, doi:10.1111/ejn.13374 (2016).

72 Jaramillo, A. A. et al. Functional role for suppression of the insular-striatal circuit in modulating interoceptive effects of alcohol. Addict Biol, doi:10.1111/adb.12551 (2017).

73 Gehrlach, D. A. et al. Aversive state processing in the posterior insular cortex. Nat Neurosci 22, 1424–1437, doi:10.1038/s41593-019-0469-1 (2019).

74 Paxinos, G. & Watson, C. The rat brain in stereotaxic coordinates. 4th edn, (Academic Press, 1998).

